# Towards new AQP4 inhibitors: ORI-TRN-002

**DOI:** 10.1101/2023.11.22.568252

**Authors:** Michael Thormann, Nadine Traube, Nasser Yehia, Roland Koestler, Gergana Galabova, Nanna MacAulay, Trine L. Toft-Bertelsen

**Author notes:** Correspondence to Trine L. Toft-Bertelsen, University of Copenhagen, Faculty of Health and Medical Sciences, Department of Neuroscience, Blegdamsvej 3, DK-2200 Copenhagen N, Denmark;.

## Abstract

Cerebral edema is a life-threatening condition that can cause permanent brain damage or death if left untreated. Existing therapies aim at mitigating the associated elevated intracranial pressure, yet they primarily alleviate pressure rather than preventing edema formation. Prophylactic anti-edema therapy necessitates novel drugs targeting edema formation. Aquaporin 4 (AQP4), an abundantly expressed water pore in mammalian glia and ependymal cells, has been proposed to be involved in cerebral edema formation. A series of novel compounds have been tested for their potential inhibitory effects on AQP4. However, selectivity, toxicity, functional inhibition, sustained therapeutic concentration, and delivery into the central nervous system are major challenges. Employing extensive DFT calculations, we identified a previously unreported thermodynamically stable tautomer of the recently identified AQP4-specific inhibitor TGN-020. This novel form, featuring a distinct hydrogen bonding pattern, served as a template for a COSMOsim-3D based virtual screen of proprietary compounds from Origenis™. The screening identified ORI-TRN-002, an electronic homologue of TGN-020, demonstrating high solubility and low protein binding. Evaluating ORI-TRN-002 on AQP4-expressing *Xenopus laevis* oocytes using a high-resolution volume recording system revealed an IC_50_ of 2.9 ± 0.6 μM, establishing it as a novel AQP4 inhibitor. ORI-TRN-002 exhibits superior solubility and free fraction limitations compared to other reported AQP4 inhibitors, suggesting its potential as a promising antiedema therapy for treating cerebral edema in the future.

## Introduction

Cerebral edema is excess accumulation of fluid in the intracellular or extracellular spaces of the brain (*1*) and prevalent in various acute brain injuries, where it significantly increases mortality independently elevates the risk of adverse outcomes. Existing therapies aim at mitigating critical consequences, such as elevated intracranial pressure, yet they primarily alleviate pressure rather than preventing edema formation, often through interventions like decompressive craniectomy where a bone flap from the skull is removed to allow the brain to expand. Water can cross cell membranes via water channels (aquaporins; AQP) and cotransport proteins (*2*). Of the large family of AQPs expressed in the majority of cell types in the mammalian body, aquaporin 4 (AQP4) is the predominant water channel in the mammalian brain (*3*) and is mainly expressed in the perivascular glial endfeet at the brain-blood interface (*4*) and the pia, in addition to the ependymal lining bordering the fluid-filled cavities of the brain, the cerebral ventricles (*5*). This distribution has led to the suggestion that AQP4 controls water fluxes into and out of the brain parenchyma and that AQP4 plays a major role in brain edema (*5-8*). With the lack of a specific inhibitor of AQP4, these studies were all based on AQP4 knock-out mice (AQP4^-/-^), which have been found to present with several molecular changes occurring secondarily to the knock-out strategy (*9-13*). To adress the exact and still not fully defined role of AQP4 in brain physiology and pathophysiology, and circumvent the side effects of AQP4 knock-out, an AQP4-specific inhibitor is a desired additional tool. The identification of molecules that inhibit specific AQP isoforms is not a simple proposition for drug discovery on account of the high level of structural conservation within the family (*13*). Several chemical structures have been identified (*14, 15*) and suggested to have AQP4 inhibtory effects. AER-270 (N-[3,5-bis (trifluoromethyl)phenyl]-5-chloro-2-hydroxybenzamide), an investigational drug that has successfully completed a Phase I study, is found to reduce central nervous system (CNS) oedema (*16, 17*). AER-271/SIM0307 (2-([3,5-Bis(trifluoromethyl) phenyl]carbamoyl)-4-chlorophenyl dihydrogen phosphate), a soluble phosphate ester prodrug of AER-270, is found to reduce swelling in two models of CNS injury complicated by cerebral edema: water intoxication and ischemic stroke modeled by middle cerebral artery occlusion (MCAO). Of note, only 20% maximal inhibition of human AQP4 was reported by AER-270 (*16*). Another drug, the 2-(nicotinamide)-1,3,4-thiadiazole (TGN-020), was found to be the most potent AQP4 inhibitor as based on *in silico* (*14*) and *in vitro* (*18*) methods. TGN-020 represents a peculiar compound of elusive origin, likely first described as an AQP4 inhibitor in a Japanese patent application from 2006: JP2006154063, later granted in the US (patent ref. US7659312B2) and Japan (JP4273235B2) but actually not claiming TGN-020 itself, rather the chemical formulation as an active compound. Here, we performed deep structural investigation of TGN-020 to elucidate the uniqueness of this compound, and to potentially generate a new starting point for the ligand-based screening and design of novel AQP4 inhibitors. Using a platform that combines artificial intelligense (AI) technologies for disease and target selection, drug design, novelty assessment, complex decision-making, and evolutionary learning (*16, 19*), we here present a soluble and low protein binding electronic homologue of TGN-020, the ORI-TRN-002 compound, which exerts inhibitory effects on AQP4-mediated water permeability as determined with heterologously expressed AQP4 in *Xenopus laevis* oocytes (*18*).

## Results

### Identification of an electronic homologue of TGN-020

To perform an unbiased deep structural investigation of TGN-020, we used fluid thermodynamics and quantum chemical calculations (DFT/COSMO-RS) of the previously identified AQP4-selective inhibitor TGN-020 (*18*) (**Fig.1A**). To identify the lowest energy tautomer, we subjected all potentially existing tautomeric forms (**Fig.1Bi-iii**) of TGN-020 to 3D structure generation, conformational analysis, and density functional theory (DFT)-energy optimization using Turbomole BP-TZVP-COSMO. The COSMO-RS phase is unlike the gas phase, but rather presents an environment of high electron density. The most stable structure of TGN-020 in our unbiased approach represents a novel tautomer of the compound (**Fig.1Biii**). This one differs from the described X-ray, IR, and NMR structure (*20*), where TGN-020 makes strong interactions with itself (X-ray and IR), and in the aprotic solvent DMSO (NMR). Notably, the carbonyl group of the amide points to the sulfur side of the thiadiazole in this compound, in agreement with our predictions for TGN-020. Compared to the X-ray structure of TGN-020 (*20*), the hydrogen bonding pattern in the new tautomer is different in the amide and thiadiazole parts (**Fig.1C-D**). The new pattern was used as a template to screen for electronic bioisosteres, i.e., molecules with similar sterics and electron distribution by using COSMOsim-3D, which is able to identify such bioisosteres with high confidence (*15, 16*) by aligning and comparing their COSMO-RS σ-surfaces. These result from quantum chemical calculations of molecules in a simulated conductor, and their histograms, the so-called σ-profiles, are widely proven to provide a very suitable and almost complete basis for the description of molecular interactions in condensed systems. COSMOsim-3D uses local σ-profiles on a spatial grid and measures intermolecular similarities based on the 3D representation of the surface polarization charge densities σ of the target and the probe molecule. The probe molecule is translated and rotated in space to maximize the sum of local σ-profile similarities between target and probe. This sum, the COSMOsim3D similarity, is a powerful descriptor of ligand similarity and allows for a good discrimination between bioisosters and random pairs. With density functional theory (DFT)-geometry optimization, which is done by moving the atoms of the molecule to get the most stable structure with the lowest possible ground state energy, we obtain both the lowest energy forms and their COSMO-RS σ-surfaces. The new TGN-020 tautomer was used as target template (**Fig.1E**) to screen Origenis proprietary compound library and corresponding optimized conformers and their COSMO surfaces using COSMOsim-3D search (*16*). We identified ORI-TRN-002 (**Fig.1F**) as the most COSMOsim-3D similar compound in our collection (**Fig.1G**). This compound recapitulates the peculiar hydrogen bonding pattern of the new form of TGN-020 almost perfectly in shape and electronic distribution although their chemical structures vary strongly due to distinct hydrogen bonding patterns.

**Figure 1.**
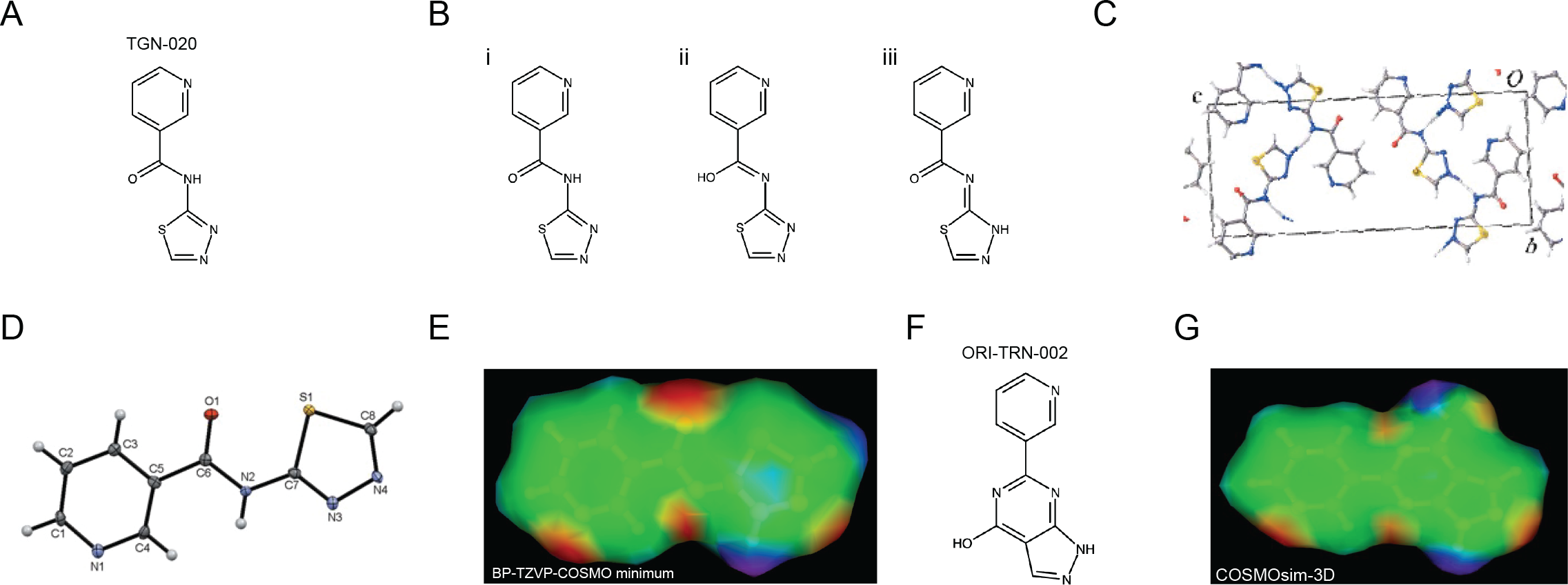
Identification of an electronic homologue of TGN-020. Commonly used chemical structure of TGN-020 **A**. Three tautomers of TGN-020 **B**. Unit cell of TGN-020 X-ray structure **C**. TGN-020 X-ray structure **D**. COSMO-RS surface of the most stable conformer of TGN-020 (**Biii**), displaying shape and hydrogen bonding pattern (red HB acceptor, blue HB donor) **E**. Chemical structure of ORI-TRN-002 **F**. COSMO-RS surface of ORI-TRN-002, displaying shape and hydrogen bonding pattern (red HB acceptor, blue HB donor), aligned with (**E**) **G**.

### ORI-TRN-002 displays drug-like properties

To evaluate potential drug-like properties of the electronic homologue ORI-TRN-002, we measured the compounds’ properties in the Origenis BRAINstorm™ platform. This platform uses affinity chromatography under standard conditions and standard calibrants to measure compound properties with high accuracy and reproducibility. It allows us to assess important physiochemical properties of the compound, i.e., human and rat serum albumin affinities (logKaHSA; logKaRSA), percentage of binding to human plasma proteins (hPPB%), alpha acid glycoprotein affinity (logKaAGP), unbound fraction in liver and hepatocytes (f_u,liver_; f_u,hep_) and the octanol-water distribution coefficient at pH=7.4 (logD), which is an appropriate descriptor of lipophilicity and indicator of solubility. (**Fig.2)**, and others to finally derive tissue-specific free fractions and tissue distribution already early in the process ensuring the best-possible optimization of compounds for their planned application. ORI-TRN-002 (**Fig.2**) displays drug-like physicochemical properties similar to TGN-020, such as aqueous solubility, which is critical to drug delivery and hydrophobicity that play key roles in drug absorption, transport, and distribution (**Fig.2**). It supersedes the solubility and free fraction limitations of the Aeromics/Simcere drug AER-270 (*21*) (**Fig.2**). ORI-TRN-002 requires no prodrug approach and serves as an ideal starting point for further optimization.

**Figure 2.**
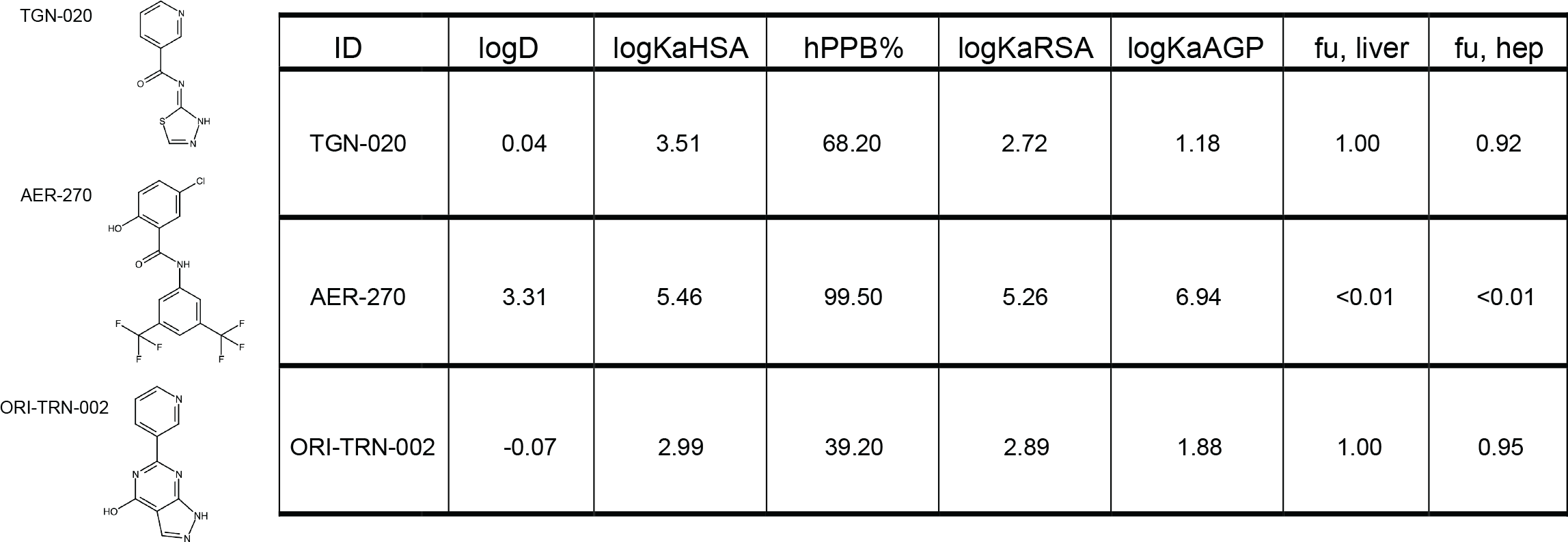
ORI-TRN-002 displays drug-like properties. The chemical structures of the three AQP4 inhibitors studied and their measured physico-chemical properties (logD at pH 7.4, log10 od molar human serum albumin affinity constant, percent human plasma protein binding, log10 of human alpha acid glycoprotein affinity constant, unbound fraction in liver, and unbound fraction in hepatocytes.

### ORI-TRN-002-mediated inhibition of AQP4

To resolve ORI-TRN-002’s inhibitory effect on AQP4, rat AQP4 (rAQP) was heterologously expressed in *Xenopus laevis* oocytes and the AQP4-mediated water permeability quantified. Following introduction of an abrupt hyposmotic challenge (-100 mOsm), we determined the osmotic water permeability of the oocytes during continuous recording of the oocyte volume with a high-resolution volume recording system. This experimental setup allows the osmotic gradient to be introduced at a faster rate than the rate of cell volume change and sufficiently fast data acquisition to obtain several data points at the initial linear part of the cell volume change, which is required to obtain the osmotic water permeability of an AQP-expressing oocyte. The native oocyte membrane has a low inherent water permeability, thus the uninjected oocytes display a negligible background (see **Fig. 3A** for summarized data on osmotic water permeabilities (compare 0.21 ± 0.13 × 10^-3^ cm sec^-1^, n = 9 for uninjected oocytes to 3.80 ± 0.33 × 10^-3^ cm sec^-1^, n = 11 for AQP4-expressing oocytes; representative volume traces shown as insets)). The oocytes were exposed to the novel drugs (or control solution) for 60 min. prior to determination of water permeability. The established AQP4 blockers, TGN-020 and AER-270 served as positive controls. Application of ORI-TRN-002 (20 μM, 60 min, 19°C) reduced the water permeability of AQP4-expressing oocytes (compare 3.43 ± 0.22 × 10^-3^ cm sec^-1^, n = 11 for non-treated AQP4-expressing oocytes to 1.39 ± 0.13 × 10^-3^ cm sec^-1^, n = 12 for AQP4-expressing oocytes incubated with ORI-TRN-002, 1.05 ± 0.15 × 10^-3^ cm sec^-1^, n = 12 for AQP4-expressing oocytes incubated with AER-270 and 0.82 ± 0.09 × 10^-3^ cm sec^-1^, n = 12 for AQP4-expressing oocytes incubated with TGN-020; representative volume traces shown as insets; **Fig.3B**). The reduced osmotic water permeability induced by ORI-TRN-002 was not by a toxic effect on the cells as verified by measuring the membrane potential of the treated oocytes (-26.2 ± 0.62 mV, n = 11 for non-treated oocytes; -24.4 ± 0.74 mV, n = 12 with ORI-TRN-002 application; **Fig.C3**). ORI-TRN-002 thus acts as a novel blocker of AQP4-dependent water permeability.

**Figure 3.**
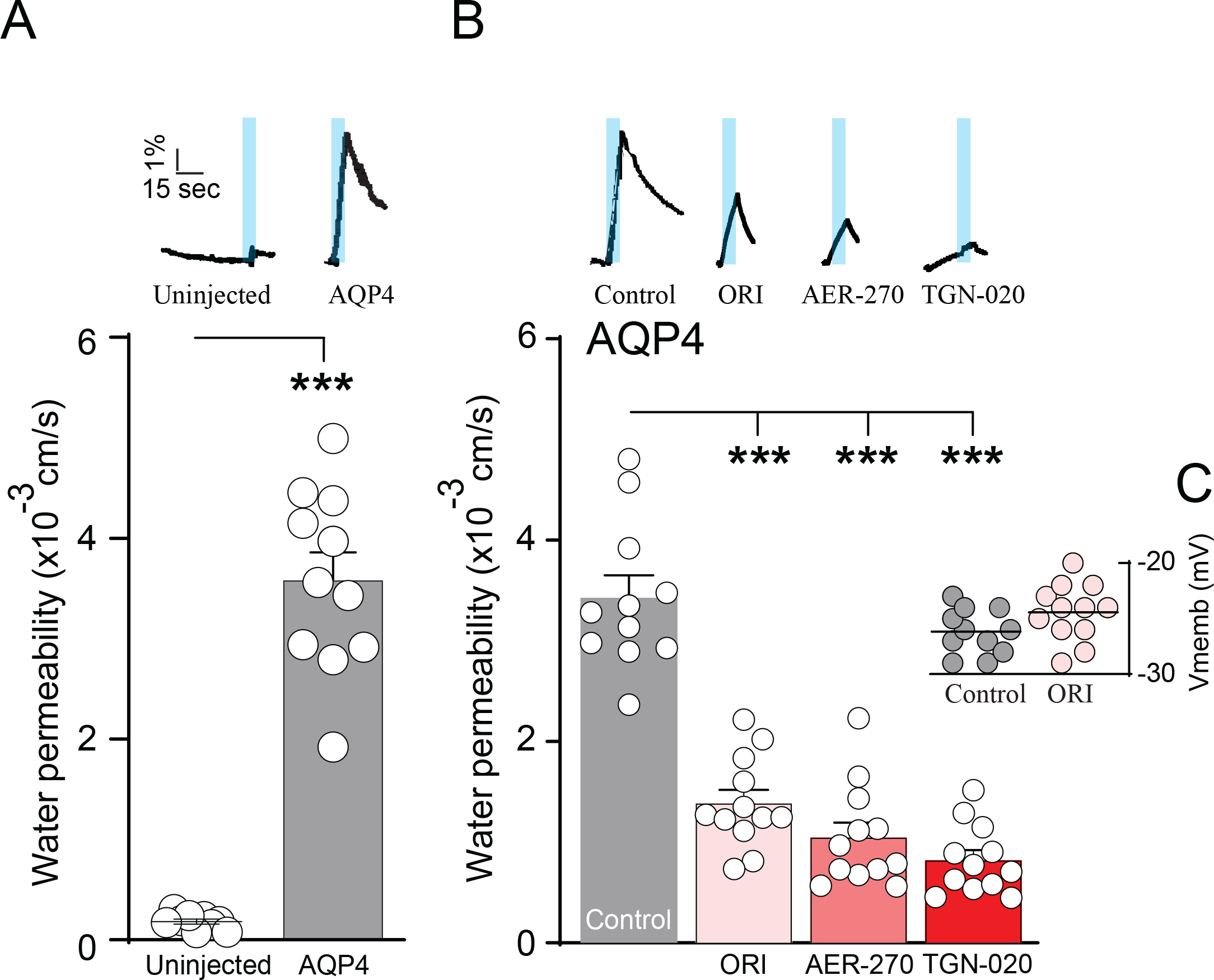
ORI-TRN-002-mediated inhibition of AQP4. Volume traces from an AQP4-expressing oocyte and an uninjected oocyte challenged with a hyposmotic gradient (indicated by a blue bar) (**A, inset**). Summarized water permeabilities from uninjected – or AQP4-expressing oocytes (n = 11) (**A**). Volume traces from an untreated AQP4-expressing oocyte or AQP4-expressing oocytes treated with ORI-TRN-002 (ORI), AER-270 or TGN-020 and challenged with a hyposmotic gradient (indicated by a blue bar) (**B, inset**). Summarized water permeabilities from oocytes expressing AQP4, either untreated or pretreated with ORI-TRN-002 (ORI), AER-270 or TGN-020 for 60 min (n = 11) (**B**). Membrane potential monitored from the AQP4-expressing oocytes in control solution or after 60 min of pretreatment with ORI-TRN-002 (ORI) (**C**) ANOVA followed by Tukeys’s multiple comparison tests or Student’s t-test were used as statistical test. ***p < 0.001.

### ORI-TRN-002 exerts an acute blocking effect on AQP4

With the observation that ORI-TRN-002 inhibits AQP4 after 60 min exposure, we subsequently demonstrated blocking efficacy within a shorter time frame. A time-control experiment verified identical magnitude of water permeabilities in the AQP4-expressing oocytes as sequentially recorded at times t=0 and t=10 min, see summarized data in **Fig.4A** (compare 3.20 ± 0.36 × 10^-3^ cm sec^-1^, n = 9 for first osmotic challenge to 3.11 ± 0.24 × 10^-3^ cm sec^-1^, n = 9 for second osmotic challenge in AQP4-expressing oocytes serving as time controls; representative volume traces shown as insets). Thus, each oocyte can be used as its own control to determine time-dependency of an inhibitor effect. Upon acute and continuous application of 20 μM ORI-TRN-002 we observed a decreased AQP4-mediated water permeability at t =10 min (compare 3.62 ± 0.24 × 10^-3^ cm sec^-1^, n = 9 for first osmotic challenge to 1.02 ± 0.17 × 10^-3^ cm sec^-1^, n = 9 for water permeability after ORI-TRN-002 application; **Fig.4A**) comparable to the inhibitory effect induced by AER-270 and TGN-020 (0.80 ± 0.11 × 10^-3^ cm sec^-1^, n = 9 after AER-270 treatment; 0.80 ± 0.08 × 10^-3^ cm sec^-1^, n = 9 with TGN-020 application; **Fig.4A**) without notable toxic effects (compare -25.6 ± 0.50 mV, n = 9 for non-treated oocytes to -26.9 ± 0.61 mV, n = 9 after ORI-TRN-002 application; **Fig.4B**). We determined the IC_50_ for ORI-TRN-002 for rAQP4 with inclusion of different ORI-TRN-002 concentrations (IC_50_ of 2.9 ± 0.6 μM, n = 6, **Fig.4C**). ORI-TRN-002 therefore appears to be a novel blocker of AQP4 with an acute effect.

**Figure 4.**
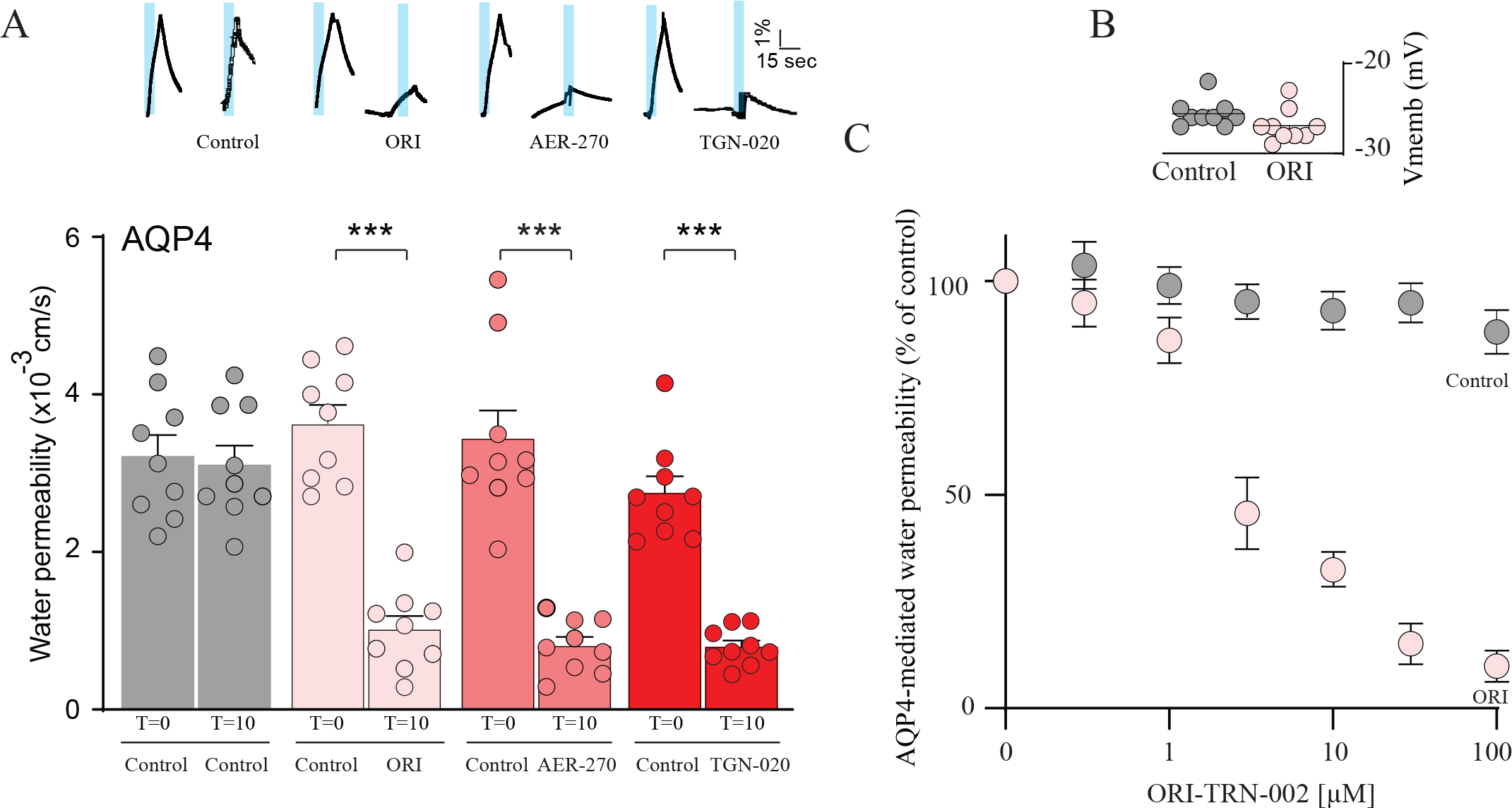
ORI-TRN-002 exerts an acute blocking effect on AQP4. A representative time control experiment from an AQP4-expressing oocyte and volume traces from AQP4-expressing oocytes treated with ORI-TRN-002 (ORI), AER-270 or TGN-020 repeatedly challenged with a hyposmotic gradient (indicated by a blue bar, t = 0 min (control), t = 10 min) (**A, inset**). Summarized water permeabilities at t = 0 min (control) and t = 10 min from oocytes expressing AQP4, either untreated or treated with ORI-TRN-002 (ORI), AER-270 or TGN-020 (n = 9) (**A**). Membrane potential monitored from the AQP4-expressing oocytes in control solution or upon treatment with ORI-TRN-002 (ORI) (**B**). AQP4-expressing oocytes exposed to different concentrations of ORI-TRN-002, data were fitted with Graphpad Prism using nonlinear regression analysis to obtain the IC_50_ from the mean of the independent regression analyses (n = 6) (**C**). ANOVA followed by Tukeys’s multiple comparison tests were used as statistical test. ***p < 0.001.

## Discussion

The current study reveals a novel AQP4 inhibitor, ORI-TRN-002, that significantly reduces the AQP4-mediated water permeability. Due to the polarized expression in the paravascular endfeet (*4*) and mediated bi-directional water flux according to the imposed osmotic gradient (*22*), AQP4 is suggested to be implicated in a range of physiological processes (*5, 23*). These findings were carried out in AQP4 knock-out mice with secondary changes to basic physiological parameters (*9-11*). Therefore, specific blockers of AQP4 are needed as supplementary tools to the knockout strategy in order to resolve a possible role for AQP4 in humans both in physiology and in cerebral edema, where the expression of AQP4 increases (*6, 24*) and edema size and formation is reduced when knocking-out (*7*) or inhibiting AQP4 (*25-27*). Current therapeutic approaches focus on alleviating critical consequences of edema, such as elevated intracranial pressure, primarily addressing pressure reduction rather than preventing edema formation. The proposed role of AQP4 in neuropathologies has heightened the importance of identifying non-toxic AQP4 activity modulators to be used for therapeutic applications. AQP drug discovery has made little progress. The small diameter of all AQP pores, together with the chemical properties of the pore-lining amino acid sidechains, means that finding drug-like compounds that can enter and block the AQPs is a challenging prospect. During the last decade, a few compounds were brought forward as potential inhibitors of AQP4; e.g., tetraethylammonium (TEA), acetazolamide, antiepileptic drugs, and bumetanide (and the derivative AqB013) (*17, 28-30*), but all of these pharmacological agents were later demonstrated to not be effective blockers of AQP4-mediated water permeability (*18, 31*). Ethoxyzolamide was reported to inhibit AQP4.M23 in oocytes (*15*), however, this compound is a carbonic anhydrase inhibitor and in patients this compound may thus induce off target effects. Other reported small molecules, although with appropriate affinity for AQP4 (*14, 15, 23, 32-34*), have yet not passed the initial phases allowing these to be enrolled in larger clinical trials. One compound that showed appropriate affinity for AQP4 and reported as a novel inhibitor by the Nakada Laboratory is TGN-020 (*14*), (*25*). Successful delivery and effect of TGN-020 to the central nervous system *in vivo* has been demonstrated (*25*) and the compound has been included in various *in vitro* and *in vivo* experimental designs (*14,18,25,33*), with a full screening on all AQP isoforms validating the isoform-specificity of TGN-020 towards AQP4 (*18*). A report from a cell-based assay could not reproduce the inhibitory results (*30*). Another compound with an appropriate affinity for AQP4 is the Aeromics/Simcere drug, AER-270, an investigational drug that was found to reduce cerebral edema. The effect was, however, only with a 20% maximal inhibition of human AQP4 (*21*), which is a little less than the inhibitory effect observed in this present study (approximately 35%). It is reasonable to suspect that the experimental approach or different species holds the explanation. AER-270 has successfully completed a Phase I study but is not yet approved by any regulatory agency. Of note, AER-270 is also a known nuclear factor (NF) kB pathway inhibitor (usually under the name IMD-0354; IKK2 Inhibitor V; N-(3,5bis(trifluoromethyl)phenyl)-5-chloro-2-hydroxybenzamide)) (*21*). A prodrug of AER-270, AER-271, selectively and effectively reduced AQP4-mediated cerebral edema and improved neurological outcome in a rodent model of transient MCAO (17), but was ineffective in reducing cerebral edema in various traumatic brain injury models (*35, 36*). This compound was still subjected to Phase I trial (NCT03804476, no data reported yet) and planned in a Phase II trial (cerebral edema prevention in acute ischemic stroke). No updates have been released yet.

From these contrasting findings, the search continues for a specific and potentially clinically relevant inhibitor of AQP4 with the same inhibitory effect no matter the test system. We reanalysed the structure of TGN-020 (*20*) and identified a novel tautomer. As evident from the COSMO-RS sigma-surface, the ORI-TRN-002 compound provides the new hydrogen bonding pattern and supports the novel tautomer as the viable target structure for AQP4. This finding contrasts the commonly used structure as reported as X-ray analysis in (*20*). Our new structure for TGN-020 is, however, not without precedence. The exo-imine tautomer of the aminothiadiazole moiety is present in the X-ray structure of 2-cyano-3-(4-(2-(2,6-dimethylphenoxy)ethoxy)-3-methoxyphenyl)-N-(5-t-butyl-(1,3,4)thiaziazol-2yl)acrylamide, which adds plausibility to our findings. ORI-TRN-002 supersedes solubility and free fraction limitations of another reported AQP4 inhibitor, the Aeromics/Simcere drug (*21*), which is a feature that requires the compound in a form of a phosphate ester prodrug due to solubility limitations. Methodological differences in executing these assays mean that attempts to replicate prior work may also result in equivocal outcomes. We here measured the osmotic water permeation through AQP4 expressed in *Xenopus laevis* oocytes using our sensitive volume measurement setup (*18*). To ascertain the osmotic water permeability of an AQP4-expressing oocyte, it is imperative to induce the osmotic gradient at a rate surpassing the cell volume change rate. Additionally, data acquisition should be rapid enough to capture multiple points during the initial linear phase of cell volume change. These fundamental technical prerequisites are at times overlooked in the study of AQP-mediated water permeability, and their absence may introduce confounding factors. We found that ORI-TRN-002 blocked AQP4 as efficiently as TGN-020 and AER-270, and the observed inhibition was evident both with pretreatment and upon acute application of ORI-TRN-002 with an IC_50_ of ∼ 3 μM. In conclusion, we here present ORI-TRN-002 as a novel and efficient inhibitor of AQP4-mediated osmotic water permeability, and with a more energetically stable form than the parent drug TGN-020 compound. AQP4 inhibition by ORI-TRN-002 is a novel approach to further address the role of AQP4 in brain water entry and exit as a complementary tool to the AQP4 knock out strategy (*37*), and could potentially be used for pharmacological treatment of pathologies involving excess AQP4-mediated fluid movements.

## Methods

### Tautomer and conformer generation

All compound structures are stored in SMILES format in an in-house database. The chemical structures were desalted and neutralized, if possible. Tautomers were generated for each compound based on the cleaned canonical SMILES using an in-house program. For each tautomer up to ten main conformers were generated using SMILES to 3D conversion and conformational sampling.

### COSMO file generation

COSMO files were generated using the TURBOMOLE program (*38, 39*) by employing geometry optimization at BP-TZVP-COSMO density functional calculation in combination with the COSMO solvation model implemented in TURBOMOLE.

### COSMO*sim3D* calculations

COSMO*sim3D* was used as described (15) to calculate the σ-surface-based similarity of the tentative bioisosteric pairs, whereby the novel tautomer of TGN-020 was compared with the lowest energy conformer/tautomer of each probe compound. The s-surfaces were extracted from the corresponding cosmo files.

### Synthesis and NMR characterisation of ORI-TRN-002

(-6-(pyridin-3-yl)-1H-pyrazolo[3,4-d]pyrimidin-4(5H)-one)

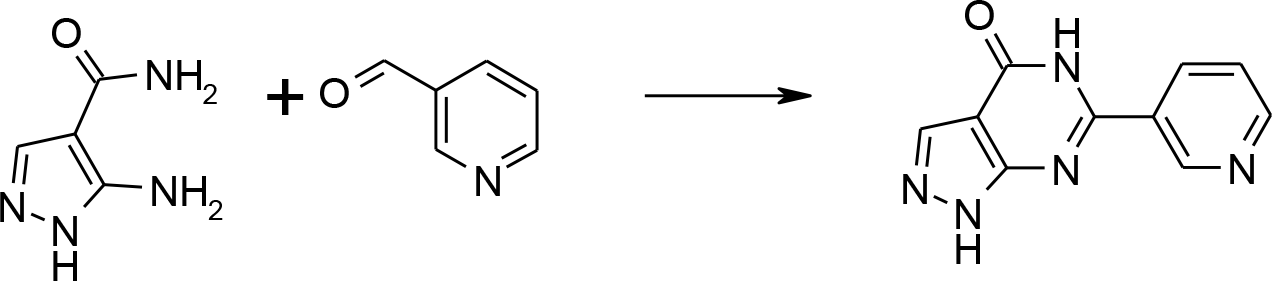

5-Amino-1H-pyrazole-4-carboxylic acid amide (5 mmol, 631 mg) and nicotinaldehyde (5 mmol, 536 mg were dissolved in HOAc (15mL). DDQ (3.75 mmol, 851 mg) was added and the mixture stirred at 140°C und microwave irritation. The mixture was filtered, the filtrate diluted with EtOAc (75mL) and filtered again. The combined solids were purified by reversed phase gradient flash chromatography. Yield 0.45 mmol, 97 mg; HPLC-MS; 98% pure, MH^+^ 214.0; ^1^H NMR (400 MHz, DMSO-d6) δ = 9.52 (d, J = 2.1 Hz, 1H), 9.14 (ddd, J = 8.3, 2.1, 1.8 Hz, 1H), 9.06 (dd, J = 5.7, 1.8 Hz, 1H), 8.26 (s, 1H), 8.18 (dd, J = 8.3, 5.7 Hz, 1H). The compound is available upon request to M.T., Origenis™.

### RNA preparation and heterologous expression in *Xenopus laevis* oocytes

The most prominent isoform of AQP4 in the brain, called M23 due to its translation initiation site at methionine at position 23 in the primary structure of AQP4 (*3*), was employed for these experiments. cDNA encoding rat AQP4 was subcloned into an oocyte expression vectors pXOOM, linearized downstream from the poly-A segment, and *in vitro* transcribed using T7 mMessage machine according to manufacturer’s instructions (Ambion, Austin, TX). cRNA was extracted with MEGAclear (Ambion, Austin, TX) and micro-injected into defolliculated *Xenopus laevis* oocytes (EcoCyte Bioscience (Germany)): 10 ng RNA was injected per oocyte and the oocytes kept in Kulori medium ((in mM): 90 NaCl, 1 KCl, 1 CaCl_2_, 1 MgCl_2_, 5 HEPES (pH 7.4)) for 3-4 days at 19°C prior to experiments.

### Volume recordings of oocytes

Oocytes were placed in an experimental recording chamber, perfused with various solutions, and volume measurements were performed as previously described (*40*). Briefly, the oocytes were placed in a small chamber with a glass bottom and perfused with solutions of interest. The volume of the oocytes was viewed from below via a long-distance objective, and micrographs were captured continuously with a high-resolution recording system (with a high signal to noise ratio) based on a CCD camera at a rate of 25 images/s (*41*). The control perfusion solution consisted of (in mM): 50 NaCl, 2 KCl, 1 MgCl_2_, 1 CaCl_2_, 10 HEPES, 100 mM mannitol (Tris buffered pH 7.4, 220 mOsm). Hyposmotic solutions were subsequently made by the removal of mannitol (Δ100 mM mannitol) with resulting osmolarity of 120 mOsm. The osmotic water permeability was determined by L_p_ = (J_v_)/(A x Δπ x V_w_), where *J*_*v*_ is the initial water flux during an osmotic challenge, *A* is the membrane surface area (nine times the apparent area due to membrane folding (*42*), Δ*π* is the osmotic challenge, and *V*_*w*_ is the partial molal volume of water (18 cm^3^/mol). Osmolarities of all solutions were verified with an accuracy of 1 mOsm with an osmometer Type 15 (Löser Messtechnik, Berlin, Germany). The compounds used were dissolved in DMSO (controls exposed to vehicle (DMSO) only) and their effect on volume changes were recorded after 10 min of treatment with the oocytes serving as own controls or after long-term application where the oocytes were incubated at 19 °C for 60 min. Determination of compound-mediated effects was carried out in a researcher-blinded fashion.

### Data presentation and statistics

All data are given as mean ± SEM. Statistical tests were performed by use of GraphPad Prism 8.4.3 (GraphPad Software Inc., La Jolla, CA, USA). Statistical significance was tested with Student’s t-test or one-way ANOVA as stated in the figure legends. *P* values < 0.05 were considered statistically significant. The number of experiments (*n*) corresponds to independent measurements from two-three different oocyte preparations. The equation y = 100/(1 + 10⋀((logIC_50_ − x) × Hill slope)) was used to fit the IC_50_ curves for each individual experiment, which are shown as summarized.

## Abbreviations

AQP: Aquaporin
AQP4: Aquaporin 4
COSMO-RS: COnductor-like Screening Model for Realistic Solvents
DFT: density functional theory
TGN-020: 2-(nicotinamide)-1,3,4-thiadiazole

## Author contributions

Conception and design of research: N.M., M.T. and T.L.T.B.; Analysis of data: N.T., N.Y., R.K., M.T. and T.L.T.B; Interpretation of results: N.M., N.T., N.Y., R.K., M.T. and T.L.T.B.; Preparation of figures: M.T. and T.L.T.B; Drafting of manuscript: M.T. and T.L.T.B.; All authors revised and approved the final version of the manuscript.

## Data availability

We confirm that all relevant data from this study are available from the first author upon request.

## References

1. V. Leinonen, R. Vanninen, T. Rauramaa, Raised intracranial pressure and brain edema. Hand Clinic 145, 25–37 (2018).

2. N. MacAulay, T. Zeuthen, Water transport between CNS compartments: contributions of aquaporins and cotransporters. Neuroscience 168, 941–956 (2010).

3. M. Amiry-Moghaddam, O. P. Ottersen, The molecular basis of water transport in the brain. Nat Rev Neurosci 4, 991–1001 (2003).

4. S. Nielsen et al., Specialized membrane domains for water transport in glial cells: highresolution immunogold cytochemistry of aquaporin-4 in rat brain. J Neurosci 17, 171–180 (1997).

5. E. A. Nagelhus, O. P. Ottersen, Physiological roles of aquaporin-4 in brain. Physiol Rev 93, 1543–1562 (2013).

6. M. C. Papadopoulos, A. S. Verkman, Aquaporin-4 and brain edema. Pediatr Nephrol 22, 778–784 (2007).

7. G. T. Manley et al., Aquaporin-4 deletion in mice reduces brain edema after acute water intoxication and ischemic stroke. Nat Med 6, 159–163 (2000).

8. G. T. Manley, D. K. Binder, M. C. Papadopoulos, A. S. Verkman, New insights into water transport and edema in the central nervous system from phenotype analysis of aquaporin-4 null mice. Neuroscience 129, 983–991 (2004).

9. N. N. Haj-Yasein et al., Glial-conditional deletion of aquaporin-4 (Aqp4) reduces blood-brain water uptake and confers barrier function on perivascular astrocyte endfeet. Proc Natl Acad Sci U S A 108, 17815–17820 (2011).

10. X. Yao, S. Hrabetova, C. Nicholson, G. T. Manley, Aquaporin-4-deficient mice have increased extracellular space without tortuosity change. J Neurosci 28, 5460–5464 (2008).

11. S. Strohschein et al., Impact of aquaporin-4 channels on K+ buffering and gap junction coupling in the hippocampus. Glia 59, 973–980 (2011).

12. X. N. Zeng et al., Aquaporin-4 deficiency down-regulates glutamate uptake and GLT-1 expression in astrocytes. Mol Cell Neurosci 34, 34–39 (2007).

13. R. M. Bill, K. Hedfalk, Aquaporins - Expression, purification and characterization. Biochim Biophys Acta Biomembr 1863, 183650 (2021).

14. V. J. Huber, M. Tsujita, T. Nakada, Identification of aquaporin 4 inhibitors using in vitro and in silico methods. Bioorg Med Chem 17, 411–417 (2009).

15. V. J. Huber, M. Tsujita, M. Yamazaki, K. Sakimura, T. Nakada, Identification of arylsulfonamides as Aquaporin 4 inhibitors. Bioorg Med Chem Lett 17, 1270–1273 (2007).

16. M. Thormann, A. Klamt, K. Wichmann, COSMOsim3D: 3D-similarity and alignment based on COSMO polarization charge densities. J Chem Inf Model 52, 2149–2156 (2012).

17. F. J. Detmers et al., Quaternary ammonium compounds as water channel blockers. Specificity, potency, and site of action. J Biol Chem 281, 14207–14214 (2006).

18. T. L. Toft-Bertelsen et al., Clearance of activity-evoked K(+) transients and associated glia cell swelling occur independently of AQP4: A study with an isoform-selective AQP4 inhibitor. Glia 69, 28–41 (2021).

19. A. Klamt, M. Thormann, K. Wichmann, P. Tosco, COSMOsar3D: molecular field analysis based on local COSMO sigma-profiles. J Chem Inf Model 52, 2157–2164 (2012).

20. M. E. Burnett, H. M. Johnston, K. N. Green, Structural characterization of the aquaporin inhibitor 2-nicotinamido-1,3,4-thiadiazole. Acta Crystallogr C Struct Chem 71, 1074–1079 (2015).

21. G. W. Farr et al., Functionalized Phenylbenzamides Inhibit Aquaporin-4 Reducing Cerebral Edema and Improving Outcome in Two Models of CNS Injury. Neuroscience 404, 484–498 (2019).

22. A. K. Meinild, D. A. Klaerke, T. Zeuthen, Bidirectional water fluxes and specificity for small hydrophilic molecules in aquaporins 0-5. J Biol Chem 273, 32446–32451 (1998).

23. M. C. Papadopoulos, A. S. Verkman, Aquaporin water channels in the nervous system. Nat Rev Neurosci 14, 265–277 (2013).

24. G. H. Tang, G. Y. Yang, Aquaporin-4: A Potential Therapeutic Target for Cerebral Edema. International Journal of Molecular Sciences 17, (2016).

25. H. Igarashi, V. J. Huber, M. Tsujita, T. Nakada, Pretreatment with a novel aquaporin 4 inhibitor, TGN-020, significantly reduces ischemic cerebral edema. Neurol Sci 32, 113–116 (2011).

26. T. Nakano et al., Prevents Brain Edema after Cerebral Ischemic Stroke by Inhibiting Aquaporin 4 Upregulation in Mice. J Stroke Cerebrovasc 27, 758–763 (2018).

27. L. Tradtrantip, B. J. Jin, X. M. Yao, M. O. Anderson, A. S. Verkman, Aquaporin-Targeted Therapeutics: State-of-the-Field. Adv Exp Med Biol 969, 239–250 (2017).

28. V. J. Huber, M. Tsujita, I. L. Kwee, T. Nakada, Inhibition of aquaporin 4 by antiepileptic drugs. Bioorg Med Chem 17, 418–424 (2009).

29. E. Migliati et al., Inhibition of aquaporin-1 and aquaporin-4 water permeability by a derivative of the loop diuretic bumetanide acting at an internal pore-occluding binding site. Mol Pharmacol 76, 105–112 (2009).

30. A. S. Verkman, A. J. Smith, P. W. Phuan, L. Tradtrantip, M. O. Anderson, The aquaporin-4 water channel as a potential drug target in neurological disorders. Expert Opin Ther Tar 21, 1161–1170 (2017).

31. M. Abir-Awan et al., Inhibitors of Mammalian Aquaporin Water Channels. Int J Mol Sci 20, (2019).

32. M. G. Mola, G. P. Nicchia, M. Svelto, D. C. Spray, A. Frigeri, Automated cell-based assay for screening of aquaporin inhibitors. Anal Chem 81, 8219–8229 (2009).

33. V. J. Huber, M. Tsujita, T. Nakada, Aquaporins in drug discovery and pharmacotherapy. Mol Aspects Med 33, 691–703 (2012).

34. F. Wang, X. C. Feng, Y. M. Li, H. Yang, T. H. Ma, Aquaporins as potential drug targets. Acta Pharmacol Sin 27, 395–401 (2006).

35. J. Wallisch et al., Effect of the Novel Aquaporin-4 Antagonist Aer-271 in Combined Tbi Plus Hemorrhagic Shock in Mice. Crit Care Med 43, (2015).

36. P. M. Kochanek et al., Operation Brain Trauma Therapy: 2016 Update. Mil Med 183, 303–312 (2018).

37. M. Eilert-Olsen et al., Deletion of aquaporin-4 changes the perivascular glial protein scaffold without disrupting the brain endothelial barrier. Glia 60, 432–440 (2012).

38. S. G. Balasubramani et al., TURBOMOLE: Modular program suite for quantum-chemical and condensed-matter simulations. J Chem Phys 152, (2020).

39. A. Schäfer, A. Klamt, D. Sattel, J. C. W. Lohrenz, F. Eckert, COSMO Implementation in TURBOMOLE:: Extension of an efficient quantum chemical code towards liquid systems. Phys Chem Chem Phys 2, 2187–2193 (2000).

40. T. Zeuthen, E. Zeuthen, N. Macaulay, Water transport by GLUT2 expressed in Xenopus laevis oocytes. J Physiol 579, 345–361 (2007).

41. T. Zeuthen, B. Belhage, E. Zeuthen, Water transport by Na+-coupled cotransporters of glucose (SGLT1) and of iodide (NIS). The dependence of substrate size studied at high resolution. J Physiol 570, 485–499 (2006).

42. G. A. Zampighi et al., A method for determining the unitary functional capacity of cloned channels and transporters expressed in Xenopus laevis oocytes. J Membr Biol 148, 65–78 (1995).

